# SPINT2 controls SARS-CoV-2 viral infection and is associated to disease severity

**DOI:** 10.1101/2020.12.28.424029

**Authors:** Carlos Ramirez Alvarez, Ashwini Kumar Sharma, Carmon Kee, Leonie Thomas, Steeve Boulant, Carl Herrmann

**Affiliations:** Health Data Science Unit, Medical Faculty Heidelberg and BioQuant, Heidelberg, Germany; Department of Infectious Diseases, Virology, Heidelberg University, Germany; Research Group “Cellular Polarity and Viral Infection”, German Cancer Research Center (DKFZ), Heidelberg, Germany

## Abstract

COVID-19 outbreak is the biggest threat to human health in recent history. Currently, there are over 1.5 million related deaths and 75 million people infected around the world (as of 22/12/2020). The identification of virulence factors which determine disease susceptibility and severity in different cell types remains an essential challenge. The serine protease *TMPRSS2* has been shown to be important for S protein priming and viral entry, however, little is known about its regulation. *SPINT2* is a member of the family of Kunitz type serine protease inhibitors and has been shown to inhibit *TMPRSS2*. Here, we explored the existence of a co-regulation between *SPINT2*/*TMPRSS2* and found a tightly regulated protease/inhibitor expression balance across tissues. We found that *SPINT2* negatively correlates with SARS-CoV-2 expression in Calu-3 and Caco-2 cell lines and was down-regulated in secretory cells from COVID-19 patients. We validated our findings using Calu-3 cell lines and observed a strong increase in viral load after *SPINT2* knockdown. Additionally, we evaluated the expression of *SPINT2* in datasets from comorbid diseases using bulk and scRNA-seq data. We observed its down-regulation in colon, kidney and liver tumors as well as in alpha pancreatic islets cells from diabetes Type 2 patients, which could have implications for the observed comorbidities in COVID-19 patients suffering from chronic diseases.

## Introduction

SARS-CoV-2 entry requires a two-step process: first, the envelope protein spike (S) binds to the viral cellular receptor Angiotensin-converting enzyme 2 (ACE2) membrane protein ^1^ and is then proteolytically activated by cellular serine proteases like TMPRSS2, TMPRSS4 and Furin ^2–4^. TMPRSS2 has been proposed as a putative drug target ^3,5,6^ and as a biomarker for COVID19 disease severity ^7,8^. Despite its central role, the regulation of *TMPRSS2* is poorly understood, although its activation by androgen response elements has been documented in normal and tumor prostate tissues ^9^.

SPINT2, a member of the Kunitz-type serine proteases inhibitors ^10^ has been shown to inhibit TMPRSS2 protease activity ^11,12^ which could have implications for COVID-19 disease. A down-regulation of *SPINT1, SPINT2* and *SERPINA1* has been reported in colorectal Caco-2 cells infected with SARS-CoV-2 ^13^, however, a clear association between *SPINT2* activation and viral permissivity has not been confirmed. To fill this gap, we evaluated the existence of coregulation between *SPINT2*/*TMPRSS2* and found common transcription factors (TF) associated with genomic loci for both genes which was also in line with a consistent correlation of the genes across cell types. This coregulation suggested a modulation of SARS-CoV-2 infection by *SPINT2*. We could corroborate a negative correlation between *SPINT2* gene expression and SARS-CoV-2 viral load in Calu-3 and Caco-2 cell lines. To validate our findings we knocked-down *SPINT2* in Calu-3 and observed an increase in the number of SARS-CoV-2 infected cells. We hypothesized that *SPINT2* levels would be lower in SARS-CoV-2 target cells from COVID-19 patients with severe symptoms which we could indeed observe in secretory cells from nasopharynx samples. This suggests that *SPINT2* can be used as a biomarker for disease susceptibility. Finally, it is known that *SPINT2* is down-regulated among different types of tumors ^14^ and we were able to corroborate this by systematically evaluating bulk- and scRNA-seq datasets which suggest a possible association to the COVID-19 comorbidity observed in cancer patients.

## Results and Discussions

### SPINT2 and TMPRSS2 are coregulated across tissues

TMPRSS2 proteolytic activity inhibition by *SPINT2* has been previously reported ^11,12^. We investigated a coregulation between *SPINT2* and *TMPRSS2*, as a similar shared regulation through the transcription factor (TF) *CDX2* has been described for *SPINT1* and *ST14 (*Matriptase) previously in enterocytes ^15^. Since *SPINT2* is also able to regulate *ST14* activity in small and large intestines ^16^ we decided to use enterocytes as a model to test this hypothesis. In order to find common TFs regulators of *SPINT2*/*TMPRSS2* we performed two independent analysis: *i*) A footprinting analysis of chromatin open regions using ATAC-seq data from Human Intestinal Organoids ^17^ identifying potential TF binding sites and *ii*) Using scRNA-seq data from ileum derived organoids ^18^ we calculated the activity of transcription factors based on the gene expression of their targets using the SCENIC algorithm ^19^. TF activities were then correlated to *SPINT2* and *TMPRSS2* gene expression. We identified common TFs inferred to be bound to the open chromatin sites in the *SPINT2* and *TMPRSS2* genomic loci (**Figure 1A**, top and bottom, respectively) and those with TF activities positively correlated to both *SPINT2* and *TMPRSS2* gene expression (**Figure 1B**, top right quadrant). Comparing these two sets of TFs, we found ten shared regulators: *ELF3, FOS, FOSL1, FOXC1, IRF1, IRF7, JUND, JUNB, ONECUT3* and *KLF4*. Interestingly, many of these regulators play a role in immune response upon infection, suggesting a possible feedback mechanism. SARS-CoV-2 infection has been shown to upregulate *FOS* expression in Huh7.5 and A549 cell lines ^20^. *IRF1* and *IRF7* are interferon regulatory factors which regulate infection responses, and have been observed to be upregulated in COVID-19 patients ^21^. *JUNB* has been found in SARS-CoV-2 infection gene expression signatures in Calu-3 and Caco-2 cell lines ^22^. *ELF3* is an important factor controlling the development of epithelium tissues ^23^ including intestinal epithelia ^24^. Based on our analysis, we depict the regulatory model in which *SPINT2* and *TMPRSS2* gene expression are coregulated by common TFs in ileum enterocytes, hence maintaining the protease/inhibitor balance (**Figure 1C**). On the other hand, down-regulation of TMPRSS2 enzymatic activity by SPINT2 possibly maintains viral load at a low level. In line with this, we observed that the expression of *SPINT2* and *TMPRSS2* are positively correlated across normal human tissues (**Figure 1D**). Interestingly, tissues which have been shown to be targets for the virus display the highest correlation between *SPINT2* and *TMPRSS2* expression. Also, at single cell resolution, we observed that both genes are specifically co-expressed in many cell types ^25^ (**Figure 1E**) which corroborates the inferred coregulation.

**Figure 1:**
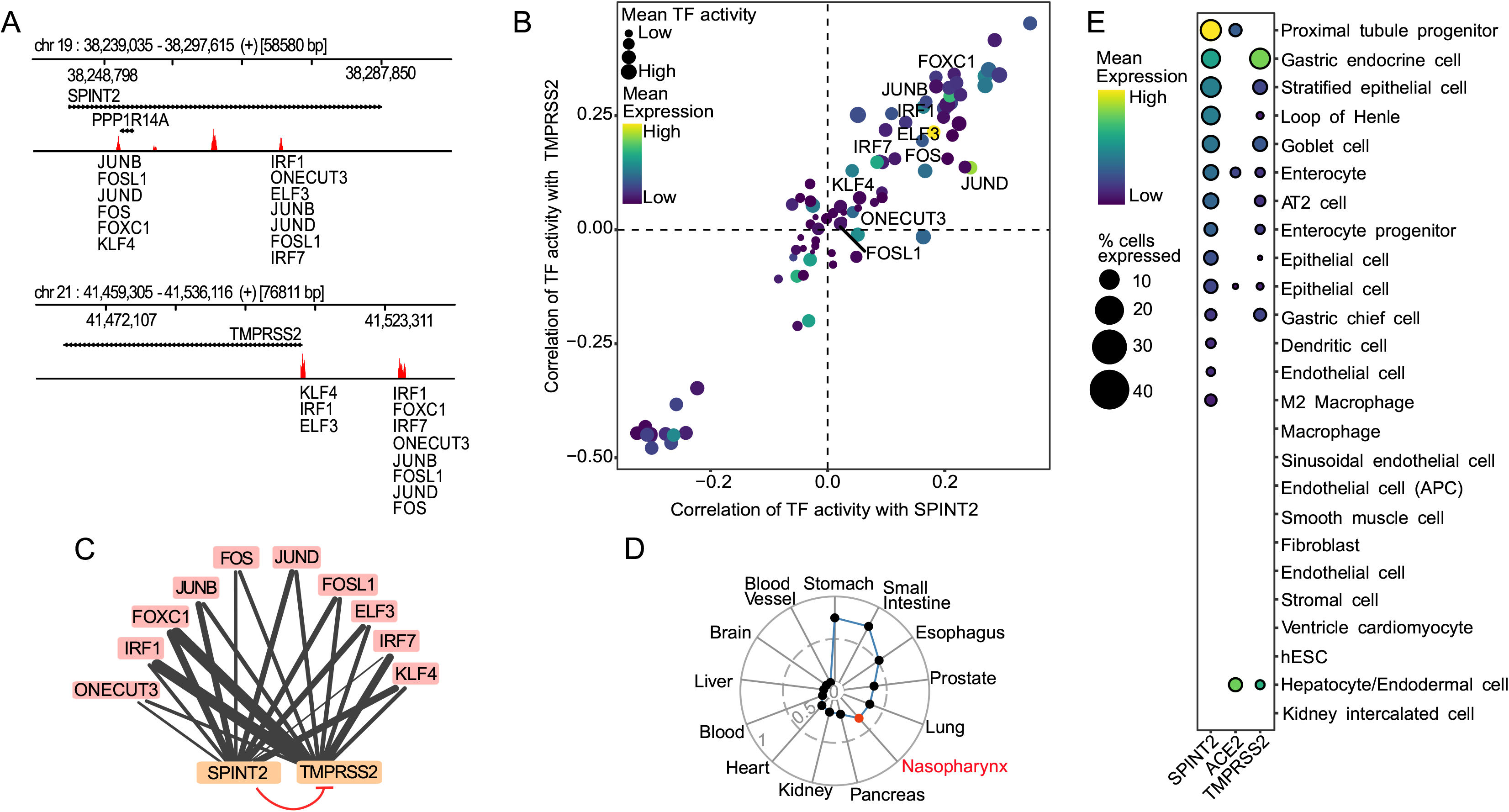
Coregulation of *SPINT2* and *TMPRSS2*. **A**. Peaks associated to open chromatin sites in regions containing the *SPINT2* and *TMPRSS2* genes (+/- 5kb of the coding regions) using ATAC-seq data from Human Intestinal Organoids. TF Binding Sites (TFBS) are scored based on the height and width of the peaks and the similarity of the sequences in the peaks to TF motifs. **B**. Correlation values of TF activities to *SPINT2* and *TMPRSS2* gene expression levels using scRNA-seq data from ileum derived organoids. Labeled TFs in (*A)* and (*B)* correspond to the intersection of inferred TFs bound to TFBS in *(A)* and those with positive correlation of TF activities to both *SPINT2* and *TMPRSS2* in *(B)*, top right quadrant. **C**. Model for *SPINT2* and *TMPRSS2* coregulation. Edges from TFs (blue) to *SPINT2* and *TMPRSS2* (salmon) represent inferred binding shown in *(A), (B)* and the edge width reflects TF binding scores. The blunt red arrow represents the inhibition of TMPRSS2 enzymatic activity by *SPINT2*. **D**. Correlation of *SPINT2* and *TMPRSS2* across tissues using bulk RNA-seq from GTEx. The additional red point labeled as nasopharynx represents the correlation of an independent dataset from nasopharynx tissue of COVID-19 patients. **E**. Gene expression of *ACE2, SPINT2* and *TMPRSS2* across cell types using HCL scRNA-seq data.

### Deriving a SARS-CoV-2 permissivity signature

Given the previous observed association between *SPINT2* and *TMPRSS2*, a major SARS-CoV-2 virulence factor ^3^, we asked whether *SPINT2* could account for differences in viral permissivity. Calu-3 and H1299 cells have been previously reported as SARS-CoV-2 permissive and non-permissive cell lines, respectively ^22^. To that end, we inferred a SARS-CoV-2 permissivity signature using the following approach: we calculated Differentially Expressed Genes (DEGs) between non-infected Calu-3 and H1299 cells (set *z* in **Figure 2A**). Then, from this list we excluded any genes that might be related to immune responses, even in mock-infected cells. In order to do so, we calculated DEGs between the infected *vs* mock-infected cells from both Calu-3 (*x*) and H1299 cell lines (*y*) and then filtered out these genes from *z* to obtain non-viral inducible genes *(i’)* (**Figure 2A**). Using this approach, we identified a set of 480 candidate genes which might contribute to infection permissivity (see **Supplementary Table 1**). We used normalized expression values for these 480 genes as input to train a Random Forest (RF) model for predicting the cumulative sum of viral gene expression in Calu-3 infected cells (**Figure 2B**). We then used the top 25% ranked genes for further analysis. *SPINT2* was found among the top ranked genes (**Figure 2B**). Of the top ranked genes, 21 corresponded to genes with functional annotations related to viral infection (**Figure 2B** inset pie chart, **Supplementary Table 1**), four have been reported to participate and/or interact directly with SARS-CoV-2 ^26^, eight corresponded to curated receptors ^27^ or ligands in the CellPhone Data Base ^28^ and 13 are cell membrane surface proteins. Importantly, by intersecting the set of genes in the permissivity signature with a reported SARS-CoV-2 viral infection transcriptional signature ^29^, we only found 3 hits (*CXCL5, LGALS3BP* and *EHF*), confirming that most of the identified genes in our permissivity signature are not viral-induced, but likely represent *a priori* susceptibility factors. We found the following Heat Shock Proteins (HSP): *HSPB1, HSPA8* and *HSPD1* to be differentially expressed. In a previous study a different HSP, *HSP90*, was observed to correlate to SARS-CoV-2 viral load in Calu-3 cells and its inhibition reduced viral infection ^22^. We also found several ribosomal proteins (*RPL9, RPL23, RPL26, RPL28, RPL38, RPS7, RPS12* and *RPS27A*) and elongation factors (*EIF3A, EIF4A2* and *EIF4B*) which could be related to viral protein translation and ER stress response ^30^. In order to confirm that the permissivity signature are not just reflecting tissue specific or immune signatures, a Pathway Enrichment Analysis (PEA) was performed using the top ranked genes. Interestingly, we found an enrichment of host-viral interactions processes, protein stabilization and Endoplasmic Reticulum (ER) trafficking pathways (**Figure 2C**).

**Table 1:**
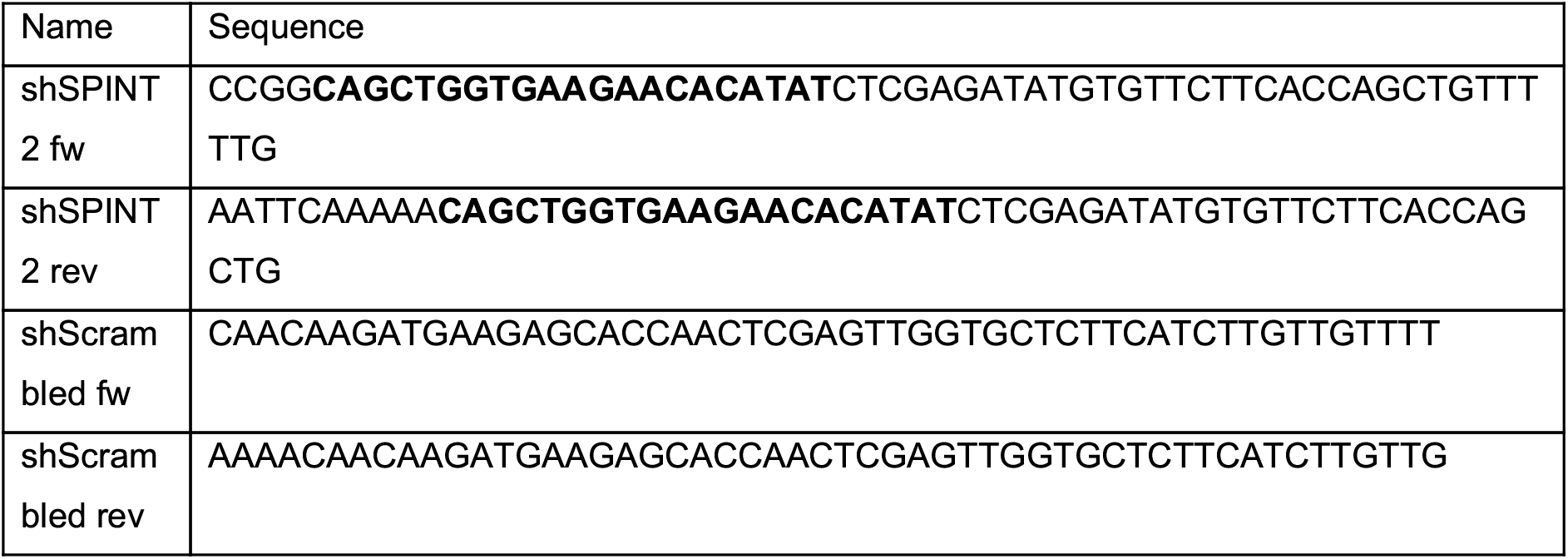
Oligonucleotides for shRNA expression. Bold characters mark the respective target sequence.

**Figure 2:**
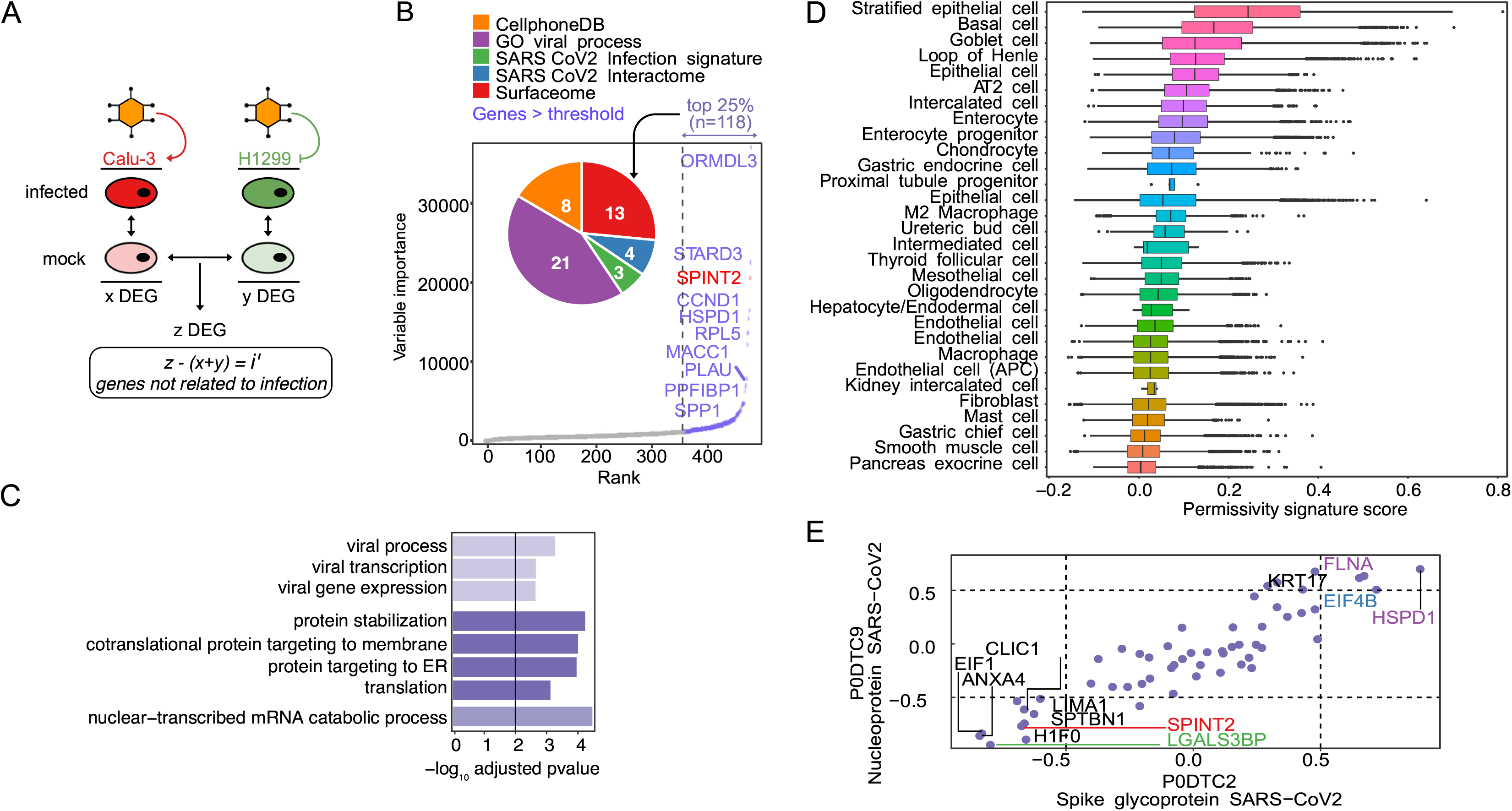
Permissivity genes are associated to viral related processes and correlated to viral expression. **A**. Permissivity signature genes (z) were found using differential gene expression analysis (DEA) between Calu-3 and H1299 permissive and non-permissive cell lines. Similarly, infection signatures for both cell lines were calculated independently (x + y) and filtered out from the permissive signature (*z*) resulting in (*i’*). **B**. Permissive genes ranked by importance in predicting the cumulative sum of gene expression of viral genes. Only the top 10 ranked genes are labeled with names. Top 25 % genes are shown in color and *SPINT2* is highlighted in red. The inset plot show the number of genes in the intersection with each geneset. See the main text for the description of each gene set. **C**. Pathway Enrichment Analysis of the top 25 % ranked genes. **D**. Ranking of cells in HCL using the permissivity scores based on the same genes as in *(B)* and *(C)*. **E**. Correlation to viral proteins (S and N genes) in the translatome data of Caco-2 cells. Gene names are coloured as in the pie chart in *(B)*.

Next, we investigated if these susceptibility genes, identified from lung-derived cell lines, are also expressed in other cell types. Therefore, we ranked the cell types in the Human Cell Landscape dataset (HCL) ^25^ based on the permissivity score derived from our top ranked genes in the permissivity signature (**Figure 2D**) and found that stratified epithelial, basal, AT2 lung cells and enterocytes were among the top-ranked cell types which correspond to cell types known to be infected by the virus ^31,32^.

In order to further refine our permissivity signature by going beyond transcriptional levels, we used protein expression levels of a previously released proteomic dataset from SARS-CoV-2 infected Caco-2 cells ^13^. We determined the Spearman correlations of the translation rates for the top ranked genes to that of the N and S viral proteins (**Figure 2E**). Some of the highly correlated genes (both negative and positive) have been previously reported to participate in viral infection processes. For example, *LGALS3BP* is a glycoprotein secreted molecule with antiviral properties observed in HIV and Hantavirus infection ^33,34^ and in the regulation of LPS induced endotoxin shock in murine models ^35^. *CLIC1* has been previously identified as a virulence factor of Merker Cell Polyomavirus (MCPyV) which is upregulated during infection and promotes the development of Merkel Cell Carcinoma ^36^. *HSPD1* has been shown to promote viral infection of HIV, HBV and Influenza viruses ^37^. Interestingly, *SPINT2* was consistently correlated to viral translation (**Figure 2E**). Furthermore, the correlation of *SPINT2* with viral gene expression is negative and this trend is consistent in both Caco-2 and Calu-3 cell lines, indicating a repressive role on SARS-CoV-2 infection (**Supplementary Figure 1A** and **B)**. Hence, these findings suggest that *SPINT2* represents a permissivity factor that negatively correlates with SARS-CoV-2 infection.

### SPINT2 knockdown increases viral load in Calu-3 cells

To experimentally validate the negative correlation of *SPINT2* expression with SARS-CoV-2 viral gene expression, we hypothesized that this gene could have a direct influence on SARS-CoV-2 infection by impairing early steps of viral entry. Hence, to test our hypothesis, we knocked-down *SPINT2* using small-hairpins RNA in the human lung carcinoma derived line Calu-3 cells. *SPINT2* expression was readily detectable in wild-type (WT) Calu-3 cells. When Calu-3 cells were transduced with a specific shRNA directed against *SPINT2, SPINT2* levels were significantly decreased compared to WT cells or cells transduced with a scrambled shRNA (**Figure 3A**). To address the impact of *SPINT2* knocked-down on the permissivity of Calu-3 cells to SARS-CoV-2, WT, scrambled and *SPINT2* knocked-down cells were infected with SARS-CoV-2 using the same multiplicity of infection (MOI). Knocked-down of *SPINT2* resulted in an almost two-fold increase in the number of cells positive for SARS-CoV-2 (**Figure 3B**-**C**). In order to test the hypothesis whether *SPINT2* could modulate viral load by regulating *TMPRSS2* expression, we monitored its fold change expression. Interestingly, *TMPRSS2* gene expression was found to be higher in *SPINT2* knocked-down cells when compared to WT or scramble cells (**Figure 3D**). Together, our results show that genetic depletion of *SPINT2* results in an increased susceptibility of Calu-3 cells to SARS-CoV-2 infection, which is in agreement with our analysis suggesting that *SPINT2* expression negatively correlates with infection.

**Figure 3:**
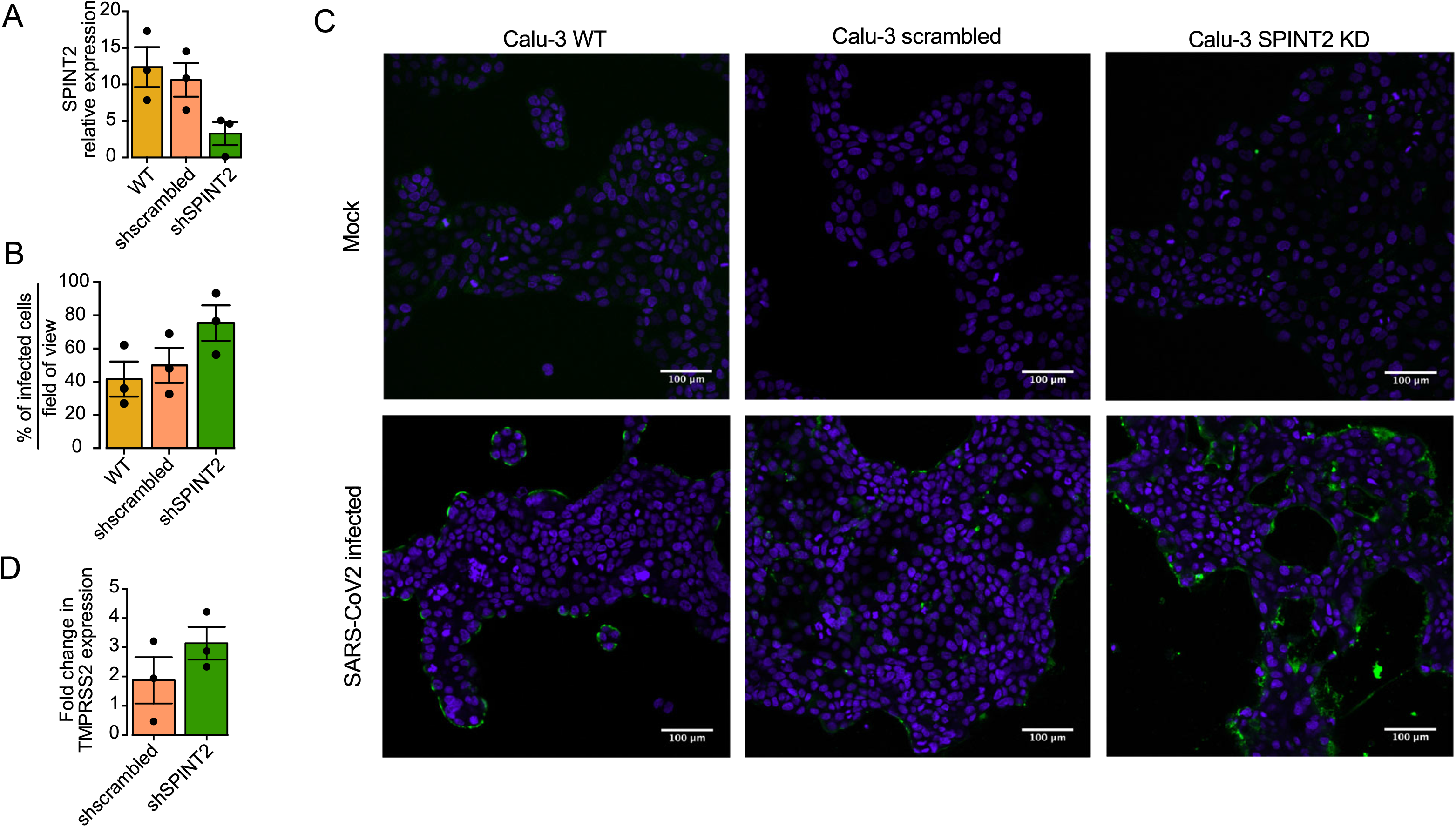
*SPINT2* knockdown in Calu-3 cell lines. **A**. Relative expression of *SPINT2* normalized to the housekeeping gene TBP in wild type Calu-3, scrambled-treated and *SPINT2* KD Calu-3 cells. **B**. Infection was detected by indirect immunofluorescence and percentage of infected cells was quantified **C**. Representative immunofluorescence images of wild type, scrambled control and *SPINT2 KD* Calu-3 cells for mock-infected and SARS-CoV-2 infected conditions. **D**. The change in *TMPRSS2* expression level was detected by qRT-PCT in WT, scrambled control and *SPINT2* KD Calu-3 cells.

### SPINT2 is negatively correlated to viral load and is down-regulated in severe COVID-19 cases

Given the observed negative correlation between *SPINT2* expression and SARS-CoV-2 infection in cell lines (**Figure 2E, Supplementary Figure 1A, B**) we next investigated if *SPINT2* expression is associated with disease severity in COVID-19 patients. We used a publicly available scRNA-seq dataset on nasopharynx swabs samples from patients with severe and mild symptoms ^38^. We correlated a list of serine proteases and inhibitors (SPRGs, **Supplementary Table 2**) to the viral RNA reads and found that *SPINT2* was the second most negatively correlated gene (**Figure 4A**). Then, we selected the cell cluster with the highest expression of *SPINT2*, which correspond to secretory cells (**Supplementary Figure 2A**) and among these cells, observed a lower *SPINT2* gene expression in cells from critical COVID19 cases compared to moderate cases (**Figure 4B**). This finding is particularly relevant since secretory cells are primary targets of viral infection ^39^. We also evaluated data on Peripheral Blood Mononuclear Cells (PBMC) from severe COVID-19 patients ^40^. In this dataset, *SPINT2* was found to be strongly expressed in Dendritic Cells (DC), plasmacytoid DC (pDC), and stem cells (SC) and eosinophils (Supplementary **Figure 2B**). Among these cells, again, we observed lower *SPINT2* expression in patients from Intensive Care Units (ICU) (**Figure 4C**). Additionally, we could also corroborate the negative correlation of *SPINT2* and viral load using bulk RNA-seq data from lung autopsies of COVID-19 deceased patients ^41^. We calculated the correlations of gene expression between *SPINT2, ACE2* and *TMPRSS2* to E, M, N and S viral genes and observed the similar negative correlation (**Supplementary Figure 2C**). Collectively, this evidence suggests that *SPINT2* expression level could be associated to COVID-19 disease severity.

**Figure 4:**
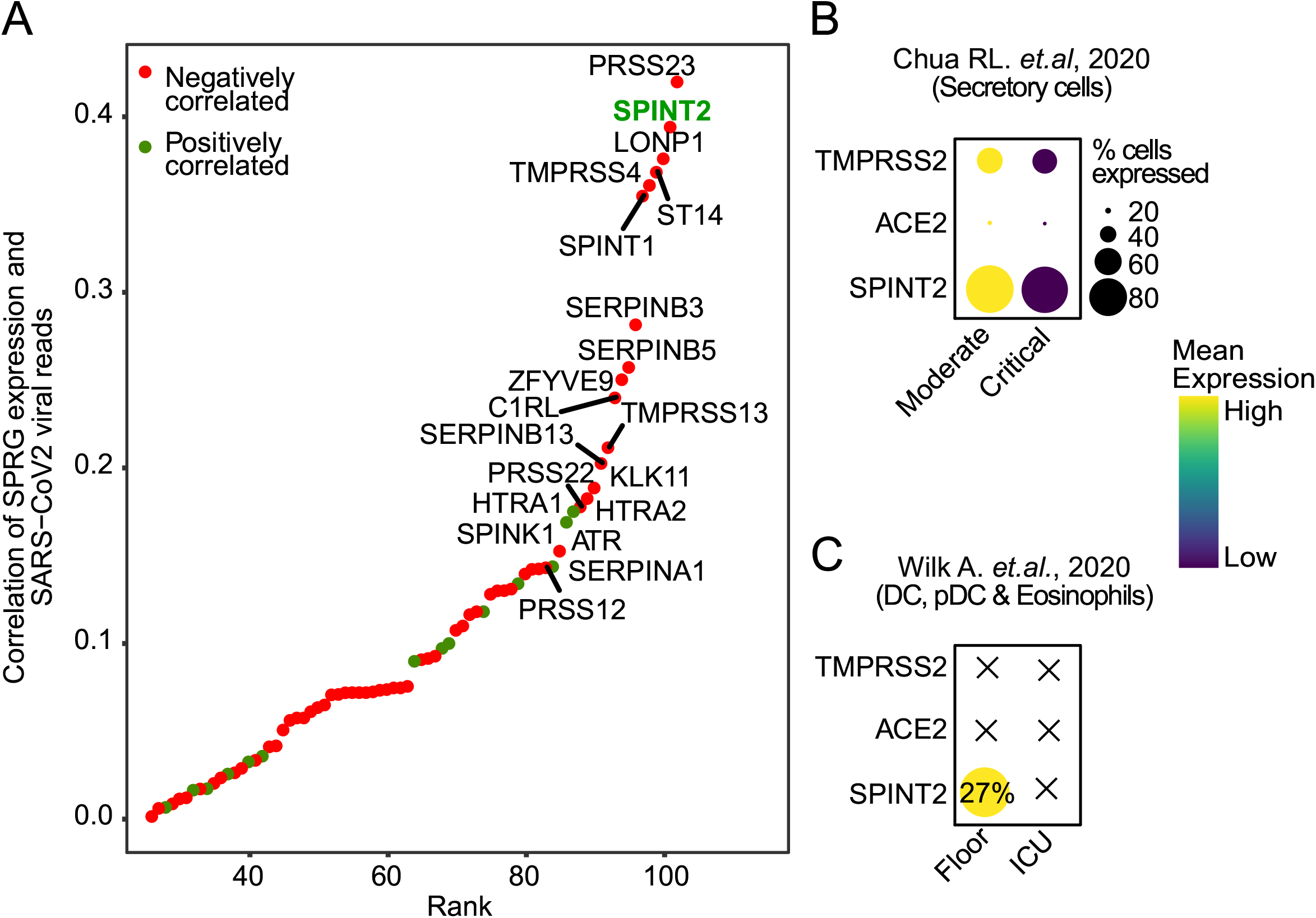
*SPINT2* expression is associated with disease severity. **A**. Spearman correlation values of SPRGs to SARS-CoV-2 viral reads in scRNA-seq of nasopharynx samples from COVID-19 patients. Top 20 most correlated SPRGs are labeled and *SPINT2* is highlighted in green. **B**. *SPINT2, ACE2* and *TMPRSS2* gene expression in severe and mild cases COVID-19 patients using datasets from Chua RL et al, 2020 **C**. Same as in (*B*) but using the Wilk A et al, 2020 data on Peripheral Blood Mononuclear Cells derived from COVID-19 patients

### SPINT2 is down-regulated in multiple tumor types and pancreatic cells from T2D patients

COVID-19 patients with previous records of chronic diseases like cancer or diabetes are considered at higher risk ^42–46^. Also, *SPINT2* gene silencing by promoter hypermethylation has been reported in multiple tumor types which promotes tumor progression ^14,47–49^. For this reason, we hypothesized that *SPINT2* down-regulation in tumor cells would increase viral infection permissivity which among others, could be one of the mechanisms behind the comorbidity observed in COVID-19 patients. We screened lung, colon, liver and hepatic tumor datasets to evaluate the differences in *SPINT2* gene expression between tumor and paired normal samples. We found statistically significant down-regulation of *SPINT2* in the kidneys and liver tumors (**Supplementary Figure 3A**). Similarly, using comparable tumor scRNA-seq datasets ^50–54^ we observed a down-regulation of *SPINT2* in colon adenocarcinoma (epithelial cells), renal clear cell carcinoma (endothelial cells) and hepatocellular carcinoma (hepatocytes) (**Figure 5**). Interestingly, we were able to detect *SPINT2* down-regulation in colorectal tumor epithelial cells at single cell level but not in bulk RNA-seq data suggesting that *SPINT2* expression might me modulated in specific cell subtypes (**Figure 5** and **Supplementary Figure 4A**). In lung adenocarcinomas, we found *SPINT2* upregulation in tumors both in TCGA and scRNA-seq data which might reflect the existence of different determinants for comorbidity in lung tissues independent of *SPINT2* modulation. We also looked at the expression of *SPINT2* in pancreatic cells from diabetes type 2 (DT2) patients ^54^. Islet cells have high *SPINT2* expression when compared to other cell types like endothelial cells (see **Supplementary Figure 4B**). We observed a strong down-regulation of *SPINT2* in alpha-cells of DT2 patients, which have been shown to be primary targets of the SARS-CoV-2 virus. This down-regulation of the virulence associated factor *SPINT2* might contribute to the comorbidity between COVD19 and DT2.

**Figure 5:**
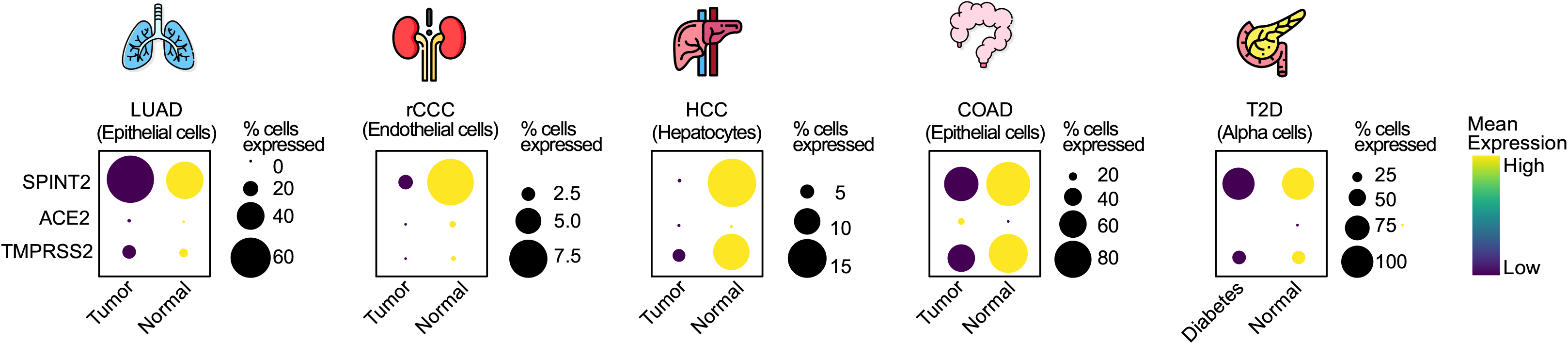
*SPINT2* is down-regulated in tumors and diabetic pancreatic cells. Dot plots showing gene expression and percentage of cells expressing *SPINT2, ACE2* and *TMPRSS2* in tumors and pancreatic alpha cells. The selected cell type for each tumor type is indicated in each case. Lung Adenocarcinoma (LUAD), renal Clear Cell Carcinoma (rCCC), Hepatocyte Cell Carcinoma (HCC), Colon Adenocarcinoma (COAD), Type 2 diabetes (T2D).

## Discussion

In this study, we describe a tight protease-inhibitor/protease balance at the gene expression level between *SPINT2* and *TMPRSS2*, a major co-receptor of SARS-CoV-2. We found Transcription Factor Binding Sites (TFBS) for ten regulators including *IRF1, IRF3, JUNB, JUND* and *ELF3* whose TF activities were found to be correlated to both *SPINT2* and *TMPRSS2* gene expression which suggests their possible role as common regulators of both genes. Interestingly, *ELF3* and *IRF7* TF activity has been found to be modulated in SARS-CoV-2 infected vs bystander enterocytes from ileum ^18^, which could point to viral load modulation mediated by *TMPRSS2* and *SPINT2* through these TFs. We show that *SPINT2* and *TMPRSS2* gene expression levels are correlated across cell types and tissues. Interestingly, known SARS-CoV-2 target tissues have high correlation values and co-expression for both genes which suggest that *SPINT2* could play a role in SARS-CoV-2 viral entry.

Currently, it is unclear what are the molecular signatures that determine viral permissivity and how they are related to disease severity. We inferred a SARS-CoV-2 permissivity signature, using differentially expressed genes between permissive and non-permissive cell lines from which we removed viral induced genes. We were able to find *SPINT2* in this permissivity signature and observed a negative correlation to SARS-CoV-2 viral load in Calu-3 cells. We also corroborated this trend at the protein level in Caco-2 cells. During the preparation of these manuscript, a study from Bojkova D et al, 2020 was published suggesting a possible role of *SPINT1, SPINT2* and *SERPINA1* in viral infection by observing the down-regulation of their protein levels in infected cells and also by evaluating the effect of Aprotinin a non-specific SP inhibitor on viral load ^6^. However, here for the first time by knocking down *SPINT2*, we provide a direct causal evidence that *SPINT2* is indeed able to modulate SARS-CoV-2 infection.

*SPINT2* inhibits *TMPRSS2* enzymatic activity through its KD1 and KD2 domains ^12^. Interestingly, we could observe an up-regulation of *TMPRSS2* mRNA expression in the *SPINT2* knocked-down Calu-3 cells. Further investigation is needed to explore this regulation at the gene expression level. Although, it has been reported previously that *SPINT2* can modulate *ST14* protein activity by regulating its shedding from the cell membrane of mouse intestinal epithelial cells ^16^ which suggest that *SPINT2* could modulate serine proteases activity through different mechanisms. Interestingly, *SPINT2* has been reported to regulate transcription of certain genes like *CDK1A* via histone methylation ^55^. Our findings suggest that *SPINT2* regulates SARS-CoV-2 viral infection through the inhibition of *TMPRSS2*, however, we cannot discard the possibility of an indirect interaction.

We found a lower expression of *SPINT2* in secretory cells from COVID-19 patients with severe symptoms ^38^. This could have implications for COVID-19 disease severity since secretory cells have been shown to be the target of SARS-CoV viral infection using organotypic human airway epithelial cultures ^39^. We found *SPINT2* in the permissivity signature from which we filtered out viral induced genes, suggesting that this gene could be used as a marker for predicting COVID-19 disease susceptibility *prior* to infection, however this needs to be further evaluated.

Serine proteases (SPs) have been reported to be abnormally regulated in diverse chronic diseases ^14,56–58^. For example, during carcinogenic development SPs influence metastasis and cancer progression ^59,60^, while in the context of diabetes they control fibrinolysis, coagulation and inflammation which in turn affects disease severity ^57^. This led us to hypothesize that shared molecular mechanisms between some chronic diseases and COVID-19 could be explained in part by the regulation of *SPINT2*. We observed *SPINT2* down-regulation in Hepatocellular Carcinoma (HCC), Colon Adenocarcinoma (COAD) and renal Clear Cell Carcinoma (rCCC) tumor cells. *SPINT2* down-regulation in liver has been reported to contribute to the development of HCC by the binding and inhibition of the serine protease *HGFA* which transforms Hepatocyte Growth Factor (*HGF*) into its active form which in turn promotes metastasis, cell growth and angiogenesis ^14,61^ and the same mechanism has been suggested for rCCC ^9^. A marked down-regulation of *SPINT2* can be observed in alpha islets pancreatic cells from diabetes patients. It has been reported that islet cells can be infected by SARS-CoV-2 which could contribute to the onset of acute diabetes ^62^. Hence, these results suggests that kidney, colon and liver tumor types as well as pancreatic islets cells from diabetic patients could be more permissive and susceptible to SARS-CoV-2 viral infection due to an imbalance of *SPINT2* gene expression, which could lead to the disruption of the protease-inhibitor/protease balance ^63^.

In conclusion, we showed for the first time that *SPINT2* is a permissivity factor that modulates SARS-CoV-2 infection. This modulation could be explained by the balance of *TMPRSS2*/*SPINT2* (serine protease/inhibitor) that we observed at the gene expression level across several tissues. We also found lower *SPINT2* gene expression in samples from COVID-19 patients with severe symptoms, hence, this gene might represent a biomarker for predicting disease severity. We also found *SPINT2* down-regulation in tumor types which could have implications for the observed comorbidities in COVID-19 patients with cancer.

## Methods

### Cell line and Viruses

Human lung adenocarcinoma cell lines Calu-3 (ATCC HTB-55) were cultured in Dulbecco’s modified Eagle’s medium (DMEM) supplemented with Glutamax (Gibco), 10% fetal bovine serum and 1% penicillin/streptomycin while Vero E6 cells (ATCC CRL 1586) were cultured in Dulbecco’s modified Eagle’s medium supplemented with 10% fetal bovine serum and 1% penicillin/streptomycin (Gibco). Calu-3 cell lines stably expressing the *SPINT2* knockdown were generated by lentiviral transduction.

### Production of lentiviral constructs expressing shRNA against SPINT2

Oligonucleotides encoding the sequence for *SPINT2* knockdown were designed from the TRC library based on Genetic Perturbation Platform (GPP) Web Portal, cloneID: TRCN0000073581 (Table 1) ^64^ Annealed oligonucleotides were ligated with the AgeI-HF and EcoRI-HF digested pLKO.1 puro vector (Add gene #8453) using the T4 DNA Ligase (New England Biolabs) and the resulting plasmids were transformed into E. coli DH5α-competent cells. Amplified plasmid DNA was purified using the NucleoBondR PC 100 kit by Marchery-Nagel following the manufacturer’s instructions.

### Lentivirus production and selection of stable cell lines

HEK293T cells (ATCC CRL-3216) were seeded on 10 cm^2^ dishes and allowed to adhere for 36 hours. The cells were transfected with 4 μg of pMD2.G (Addgene #12259), 4 μg of psPAX2 (Addgene #12260) and 8 μg of purified pLKO.1 plasmid containing the shRNA constructs upon reaching 70% confluency. Cell supernatant containing lentivirus was harvested 72 h post-transfection, filtered through a 45 μM Millex HA-filter (Merck Millipore) and purified by ultracentrifugation at 27,000x g for 90min. 2×10^5^ Calu-3 cells were seeded onto collagen coated 6-well plates 24 h prior to transduction. Cell medium was replaced with 3 mL medium containing 20 μL of the purified lentivirus and 3μl polybrene transfection reagent (Merck Millipore). Medium was supplemented with 10 µg/mL puromycin for selection of successfully transduced cells two to three days after transduction.

### SARS-CoV-2 viral infection

The SARS-CoV-2 isolate used in the experiments was obtained from the swab of a SARS-CoV-2 positive patient from the Heidelberg University Hospital. The virus was isolated and propagated in Vero E6 cells. All SARS-CoV-2 infections were performed with a multiplicity of infection of 0.04 as determined in Vero E6 cells. Prior to infection, culture media was removed and virus was added to cells and incubated for 1 hour at 37°C. Fresh media was added back to the cells upon virus removal.

### RNA isolation, cDNA synthesis and qPCR

Cells were harvested 24 hours post infection for RNA isolation using RNAeasy RNA extraction kit (Qiagen) as per manufacturer’s instructions. Complementary DNA was synthesized using iSCRIPT reverse transcriptase (BioRad) from 250 ng of total RNA per 20µL reaction according to the manufacturer’s instructions. Quantitative RT-PCR assay was performed using iTaq SYBR green (BioRad) as per manufacturer’s instructions. The expression of target genes was normalized to endogenous control *TBP*. Primer sequences are as below:

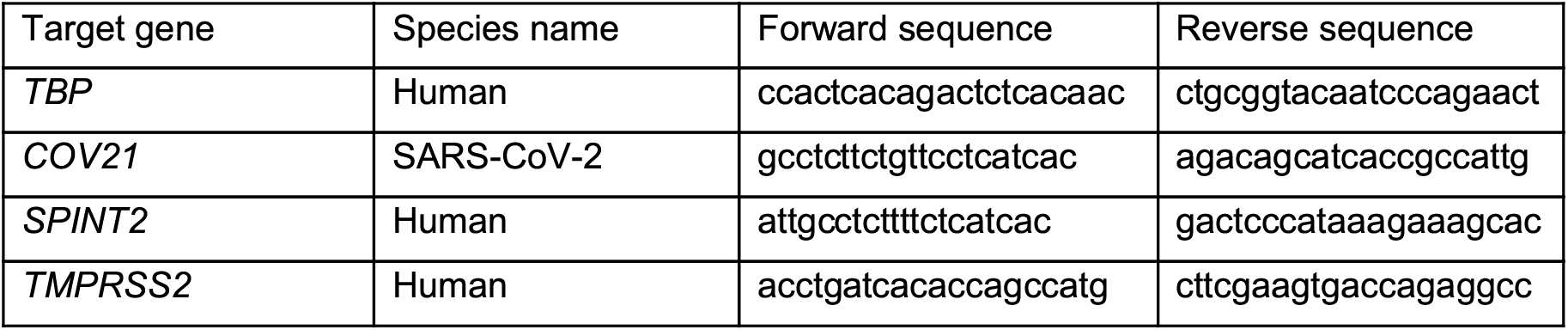

The fold change in SARS-CoV-2 genome copy number was calculated using input as a reference. Input samples were harvested directly post-infection and accounted for the basal viral genome copy number detected due to viruses attaching to the cell membrane.

### Indirect Immunofluorescence Assay

Cells were seeded on iBIDI glass bottom 8-well chamber slides which are previously coated with 2.5% human collagen in water for at least 1 hour. Cells were fixed in 4% paraformaldehyde (PFA) for 20 mins at room temperature (RT) 24 hours post infection. Cells were washed in 1X PBS and permeabilized in 0.5% Triton-X for 15 mins at RT. 30 minutes of blocking were carried out using 3% BSA-PBS at RT. Mouse monoclonal antibody against SARS-CoV-2 Nucleocapsid (NC) protein (Sino biologicals MM05) as primary antibody was diluted in 1% BSA-phosphate-buffered saline (PBS) and incubated for 1h at RT. Cells were washed with 1X PBS three times and incubated with secondary antibodies conjugated with AF488 (Molecular Probes) and DAPI for 30-45 mins at RT. Cells were washed in 1X PBS three times and maintained in PBS. Cells were imaged on a Nikon/Andor Spinning Disc Confocal microscope to quantify the number of infected cells relative to the number of nuclei.

### Statistics and computational analyses and statistics

In order to quantify infected cells from indirect immunofluorescent stained samples, ilastik 1.2.0 was used on DAPI images to generate a mask representing each nucleus as an individual object. These masks were used on CellProfiler 3.1.9 to measure the intensity of the conjugated secondary antibodies in each nucleus. A threshold was set based on the basal fluorescence of non infected samples, and all nuclei with a higher fluorescence were considered infected cells.

#### Calu-3 and 1299 cells preprocessing

For Calu-3 cells we filter out cells with an extremely high number of detected genes (>50,000) which probably corresponds to doublets. In H1299, since few cells were detected to be infected, because this line is non-permissive, in order to obtain DEGs we defined infected cells as those with cumulative sum of viral genes expression >0.

### Assessing non-viral induced permissivity signatures

As we wanted to differentiate between permissivity and infection signatures, we first looked for differentially expressed genes in SARS-CoV-2 permissive *vs* non-permissive cell lines and then we removed all the genes which were up- or down-regulated during infection (**Figure 2A**). We performed Differential Expression Analysis using Seurat ^65^ (DEA) between Calu-3 and H1299 cells in non-infected mock cells at 4 hours of culture (*z*). Then, we obtained DEGs of Calu-3 infected *vs* mock at 12 hours post infection (*x*); we did the same with H1299 infected cells *vs* mock-infected cells at 4 hpi (*y*) In all DEA we set a Log Fold Change (FC)=0.25 threshold. Finally, we removed these infection signatures from the DEGs of Calu-3 *vs* H1299 to obtain the permissivity gene signature (*i’*).

### Ranking genes using RF and Pathway Enrichment Analysis

A Random Forest (RF) regression analysis was performed using the normalized gene expression of the permissivity signature to predict the cumulative sum of the expression of viral genes in Calu-3 cells at 12 hpi. We trained the RF using a random subsample of 75% and tested the results with the remaining set. Next, we estimated the feature importance for each of the permissivity signature genes and performed enrichment analysis using enrichR ^66^ on the top 25% ranked genes.

### Scoring permissivity signatures

For the scoring of cells based on the permissivity signature among cell types in the HCL dataset, we used the top 25% RF ranked genes and applied the AddModuleScore function of Seurat setting *nbin*=100.

### SPINT2 expression correlation to viral gene expression

For the translatome correlation analysis, the summed intensity normalized values were used as provided in the study ^13^. In order to compute the correlations of SPRGs (**Supplementary Table 2**) to the viral reads in the scRNA-seq data from Chua RL et al, 2020 the raw count matrices were extracted from the Seurat object provided by the authors, splitted by sample and then imputed using scimpute ^67^ with the following parameters: drop_thr=0.5 and Kcluster equal to the number of annotated cell types in each matrix. The imputed matrices were then merged and log2 normalized. Finally, correlations were performed restricted to infected cells (viral read counts>0). In the bulk RNA-seq data from deceased COVID-19 patients log2 RPM of normalized counts are used. In both cases correlation to viral genes were carried out using spearman coefficients.

### scRNA-seq data preprocessing

In order to have a standardized workflow for the processing of scRNA-seq data we used SCT normalization using the Seurat workflow for every dataset except for Human Cell Landscape data where log2 normalization and scaling were performed since this dataset is large and using SCT was unpractical. HCC data were downloaded from GEO (GSE149614) and reprocessed. We used the Louvain method implemented in Seurat for community detection and clusters were identified by using tissue markers. We used the markers used to characterize cell types from an independent scRNA-seq human liver atlas ^68^ and using these markers identified clusters of epithelial, endothelial, hepatocytes, Kupffer and NK cells (**Supplementary Figure 3B**). For kidney, colon, prostate tumors, pancreatic cells from T2D, PBMC and Airways epithelium from SARS-CoV-2 patients’s datasets and the annotations were used as provided in the corresponding publications (see data availability section).

### SPINT2 expression in TCGA tumors data

TPM normalized counts from tumor samples in TCGA were downloaded (https://www.cancer.gov/tcga). Our analysis was restricted to tissue and sample matched tumor and normal samples only. Difference in average expression was estimated using Wilcoxon Test with Holm correction.

### Inference of Transcription Binding Sites

A footprinting analysis was carried out using the TOBIAS pipeline ^69^ with a default parameters setting of MACS –nomodel –shift −100 –extsize 200 –broad. Then, we extracted the inferred Transcription Factor Binding Sites (TFBS) for those TF with activities found to be positively correlated to both *SPINT2* and *TMPRSS2* using the single cell RNA-seq data. TFBS were visualised using the PlotTracks TOBIAS function and the network was built in Cytoscape ^70^. Edges in the network represent TF binding scores.

## Data and script availability

For this work we used the following datasets available in public repositories: scRNA-seq profiles of Calu-3 and H1299 cell lines ^22^; scRNA-seq from ileum derived organoids ^18^; translatome and proteome quantifications of Caco-2 cell line ^13^; bulk RNA-seq profiles of lung samples of COVID-19 deceased patient autopsies ^44^; Peripheral Blood Mononuclear Cells ^40^; Nasopharynx samples from COVID-19 patients ^38^; Human Cell Landscape ^25^; The Cancer Genome Atlas. Tumor scRNA-seq datasets: Human Hepatocellular Carcinoma ^51^, renal Clear Cell Carcinoma ^50^, Lung Adenocarcinoma ^53^, Colon Adenocarcinoma ^52^, pancreatic cells from Diabetes Type 2 patients ^54^; ATAC-seq data from Human intestinal organoids ^17^.

All the codes for the data processing and analysis are provided in the following GitHub repository: https://github.com/hdsu-bioquant/covid19-comorbidity.

## Supporting information

Supplementary table1 Permissivity signature with annotations

Supplementary table 2 Serine proteases related genes

## Acknowledgments

CRA acknowledges funding through the Deutsche Forschungsgemeinschaft (DFG, German Research Foundation) – Project-ID 272983813 – TRR 179. SB and CH acknowledge funding through the DFG program “Identification of the molecular origins of comorbidity in COVID-19 patients” (Project-ID 458633366).

## Figures captions

**Supplementary Figure 1.**
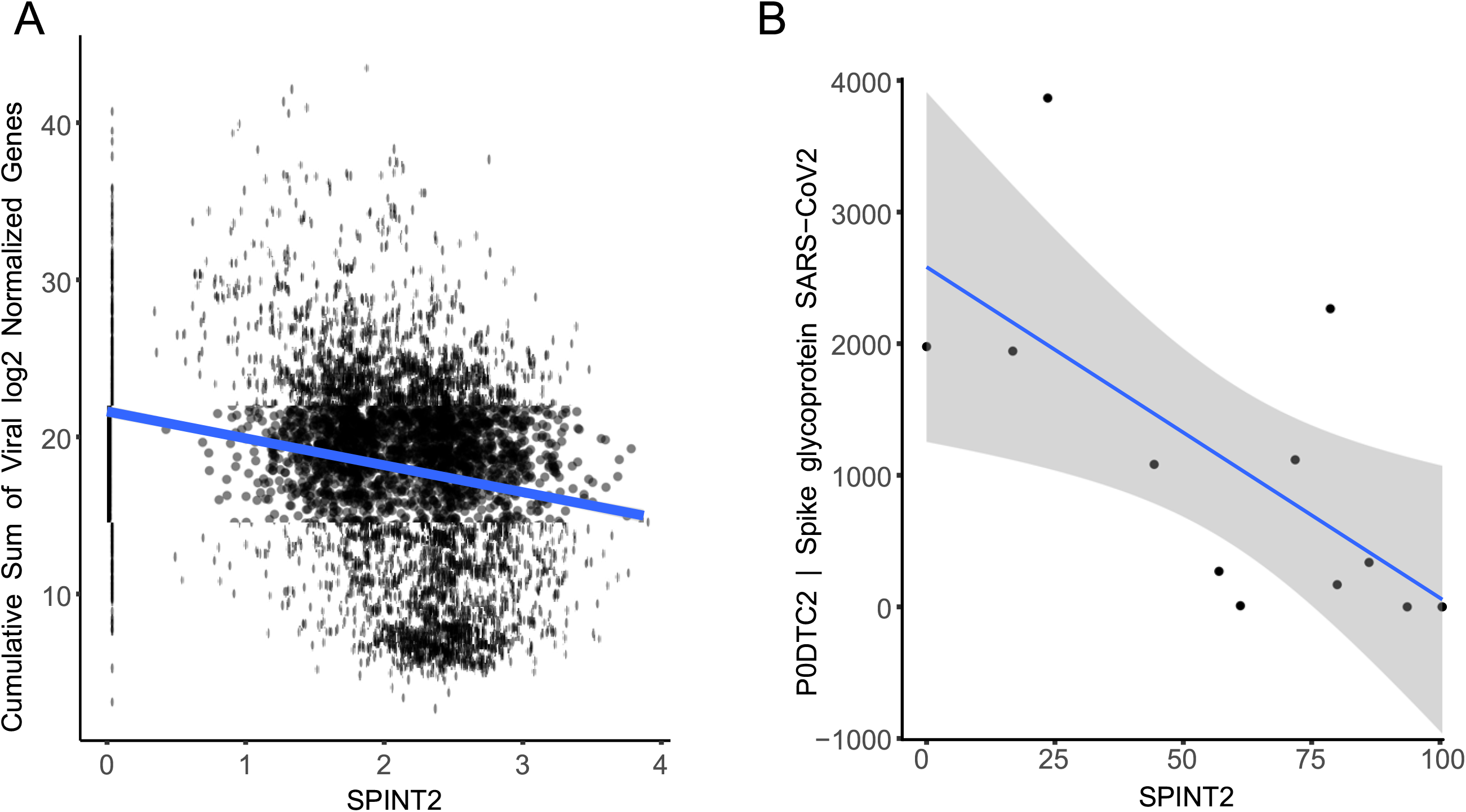
SPINT2 is negatively correlated to SARS-CoV-2 viral expression. **A**. Correlation of *SPINT2* to the cumulative sum of normalized expression values of SARS-CoV-2 genes in Calu-3 cells. **B**. Correlation of *SPINT2* to S viral protein translation rates in Caco-2 cells.

**Supplementary Figure 2.**
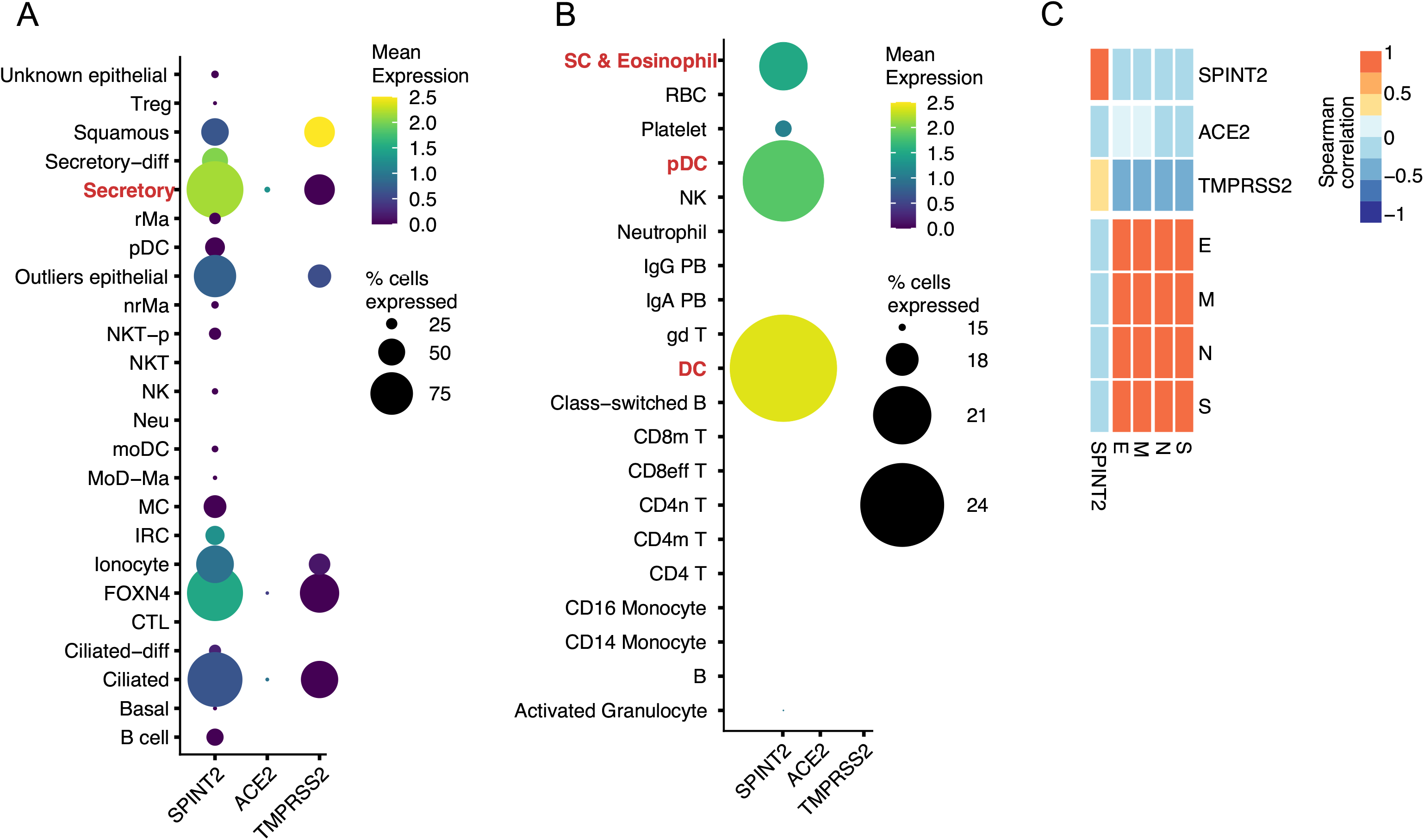
*SPINT2* expression in patients’s datasets. *ACE2, SPINT2* and *TMPRSS2* expression in Chua RL et al, 2020 (**A**) and Blish C et al, 2020 (**B**) scRNA-seq datasets. *SPINT2* expressing cell types shown in the main text are highlighted in red for both datasets. **C**. Correlation of *ACE2, SPINT2* and *TMPRSS2* to viral proteins using bulk RNA-seq from COVID-19 deceased patients in Desai N et al, 2020.

**Supplementary Figure 3.**
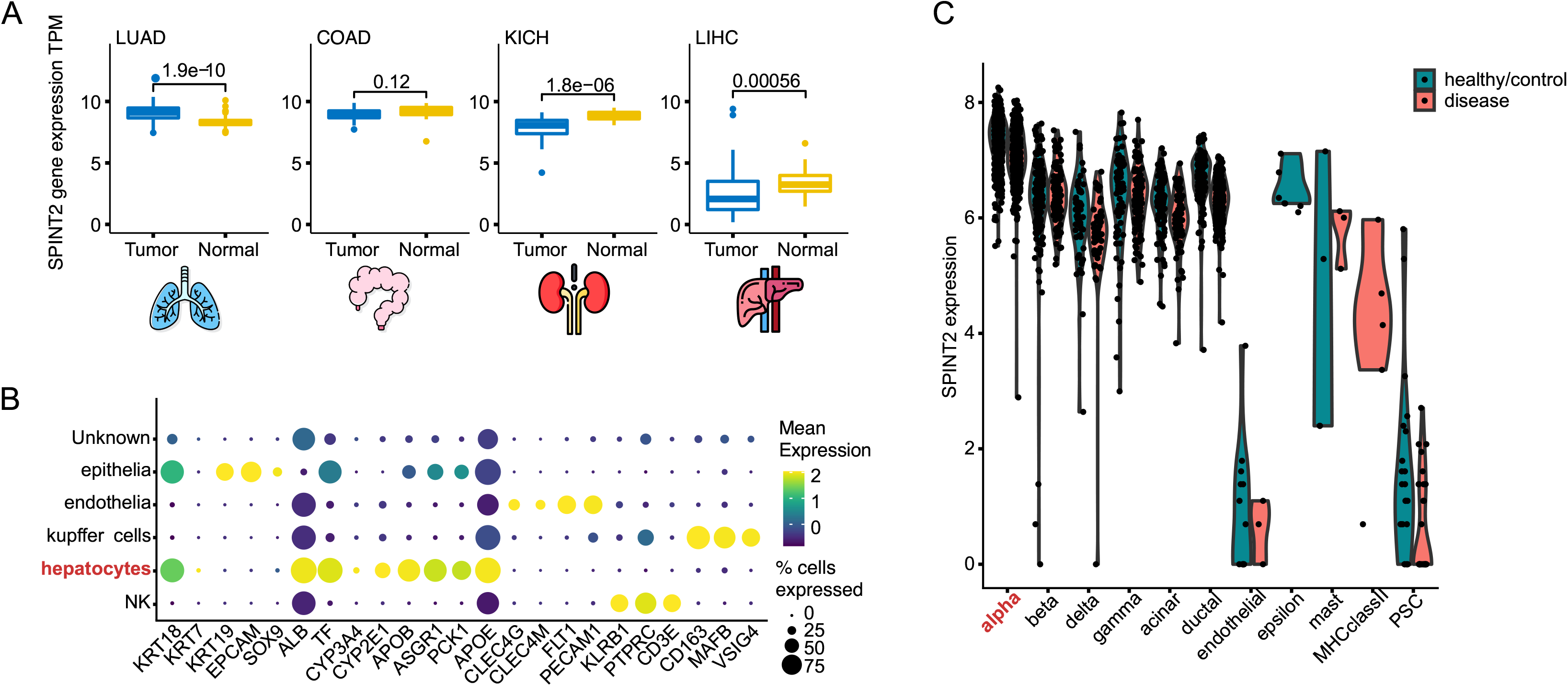
Analysis of SARS-CoV-2 associated comorbidity. **A**. Normalized gene expression (TPM) of *SPINT2* in different cancers. P-values from Wilcoxon Test. **B**. Profiling of liver cells from hepatic tumors using cell type markers as described in Aizarani N et al, 2019, see methods. **C**. *SPINT2* gene expression in pancreatic cells from diabetic patients. Hepatocytes and pancreatic alpha cells clusters highlighted in red represent cells for which gene expression profiles are shown in *Figure 5*.

